# Physiological and molecular mechanism of tolerance of two maize genotypes under multiple abiotic stresses

**DOI:** 10.1101/2021.11.11.468230

**Authors:** Suphia Rafique

**Affiliations:** Department of Biotechnology, Faculty of Chemical and Life Sciences, Jamia Hamdard, New Delhi-110062. India

**Keywords:** Multiple stresses, 2-DE, Inbred, Maize, drought x low-N stress, waterlogging x low-N stress, Tolerance vs. Susceptibility, Combined stresses

## Abstract

Abiotic stresses are the major threat to crops regardless of their nature, duration, and frequency, their occurrence either singly, and or combination is deleterious for the plant growth and development. Maize is most important crop largely grown in tropical region in summer rainy season, often face a stress combination of drought and waterlogging. We previously showed under multiple stresses up-regulated leaf proteins of maize plants were involved to enhance the tolerance mechanism of tolerant genotype. Whereas, in susceptible genotypes up-regulated proteins ameliorate to survive the stressful condition. Further to understand the response of roots proteome under multiple stresses was determined using the 2DE technique. The results of the root proteome show the up-regulated proteins of CML49 genotype (tolerant) are involved in enhancing the N content, cell wall remodeling, and acclimatization during the stresses. Up-regulated proteins of CML100 genotype (sensitive) are stressed marker of roots primary and secondary metabolism. However, the root proteome of both genotypes correlates with the leaf proteome (previous). Therefore, the present study and our previous results provide comprehensive insight into the molecular mechanisms of tolerance in multiple abiotic stresses of maize plants.

## 1. Introduction

The plants responses to multiple abiotic stresses are unique and shared. The plant responses to stress are dependent on the tissue or organ affected by the stress. The roots are the key organs that perceived stress signals and lead to brought changes at the cellular and molecular levels. Therecent climate change predictions model **(IPCC, 2014)** has predicted higher intensity of abioticstresses which are now becoming more frequent and usually results simultaneously or consecutively. They are more detrimental to crop growth, development, and production. The effect of combined stresses on crops either additive or synergistic that depends on the nature of interactions between the stress factors (**Mittler, 2006; Atkinson et al. 2013; Prasch and Sonnewald, 2013)**. Perhaps, plants are capable to be fitted their responses that may be unique and cannot be extrapolated from the response of the plant to each individual stress **(Rizhsky et al. 2004)**. Maize is the most important cereal crop that is grown at the wide geographical ranges of latitude and longitude of the world. In South Asia, particularly in tropical and subtropical environment maize is a major crop largely cultivated in the summer rainy season, often face a stress combination of drought and waterlogging due to irregular monsoon rains in the region (**Zaidi et al. 2008**). These two stress factors reveal the secondary stress effect of low nitrogen availability. N-uptake affected because of water deficit, while, in waterlogging leaching and denitrification of soil nitrogen (**Rathore et al. 1996**). Plants’ responses to combined abiotic stresses are a complex mechanism that brought changes at the transcriptome, proteome, and metabolome of the organism. The changes in cellular metabolism and recent advances in ‘Omics’ technologies have led to new insights into understanding the abiotic stress response of plants. Several workers compared the stress factor at proteome level in different treatments singly or in combined stresses **(Peng et al. 2009; Rollins et al. 2013; Oh and Komatsu, 2015)**. So far, proteomics approaches have broadened our understanding of plant stress response and for improving many physiological traits and in shaping the novel phenotype **(Kosova et al. 2018)**. In our previous work, we have shown under multiple stresses the upregulated proteins of maize leaf were involved to enhance the tolerance mechanism of the tolerant plants whereas, in susceptible plants expressed proteins help to survive the stressful condition. The present work was undertaken in continuation and roots proteome of tolerant and sensitive maize plants were compared in multiple stresses applied concurrently to understand the complex tolerance mechanism at the molecular level. Besides, the physiological analyses of the two inbred maize plants also completed to understand the differential adaptability of the two genotypes to multiple stresses.

## 2. Materials and Methods

### Plant Material and Growth conditions

In the present work two maize inbred (CML49 & CML100) genotypes were selected that showed different adaptability to various abiotic stresses applied simultaneously. The selected two genotypes (CML49 and CML100) with 80 pots each were grown in natural conditions in greenhouse up to 30 days. Thirty days after sowing (DAS) were subjected to multiple abiotic stress treatment, first 40 pots of each genotype were exposed to a drought (drought x low-N) for 10 days, while 40 control pots were supplied with full nutrient and water. After re-watering for two days normally, the same plants were exposed to waterlogging (waterlogging x low-N) for up to 7 days, to maintain the water level 2-3cm above the soil surface of the pots plants were watered day and night. The roots samples from 3-stressed and 3-control replicates plants of each genotype were kept on the last day of combined stresses for physio-biochemical analysis and extracting proteins. Roots samples were quickly frozen in liquid nitrogen after removing from the plant and then kept at −80° C for further analysis.

### Determination of Morphological parameters

To measure fresh/dry shoot and roots weights, seedlings were pulled out of the soil and roots carefully washed to remove soil particles and roots fresh weight were measured. For fresh shoot weight, the plants were cut at the stem base and measured fresh weight on electronic balance. Then stem and roots oven dried (75°C) for 48 h, for dry weight measurements by means of electronic balance. Total dry weight was calculated by adding the dry weight of shoot and root.

### Determination of Physiological parameters

Total protein was extracted by homogenizing the roots, first in liquid nitrogen, then in 4 volumes of 125 mM Tris-HCl buffer, pH 8.8, 1% (w/v) SDS, 10% (w/v) glycerol, and 1 mM PMSF. The homogenate was centrifuged for 10 min at 15000g at 4°C, and the protein content of the supernatant was determined using Bio-Rad Protein Assays Dye Reagent Concentrate and bovine serum albumin as standard. Chlorophyll was extracted by homogenizing the leaf, using mortar and pastel in 4 volumes of 80% (v/v) precooled acetone. The homogenate was centrifuged for 20 min at 12000g at 4 °C, and the chlorophyll content of the supernatant was measured at 663 and 647 nm in a spectrophotometer (Hitachi U2910). In drought x low-N stress after one week, leaf samples were collected from the second leaf for leaf RWC determinations. The relative water content (RWC) was calculated by using the formula as: RWC% = FW-DW/TW-DW x 100%.

### Determination of Biochemical parameters

In vivo assay of NR activity of roots was estimated by following the procedure, **Hageman and Hucklesby (1971)**. The roots were cut and placed in ice-cold incubation medium containing 3.0 ml of 0.2 M potassium phosphate buffer (pH 6.8), and 3.0 ml of 0.4 M KNO3 solution. The tubes were evacuated, with a vacuum pump and then incubated in a water bath at 33°C for one hour under dark conditions. At the end of incubation period, tubes were placed in boiling water bath for 5 min to stop the enzyme activity and the complete leaching of the nitrite in the medium. The calibration curve was prepared using sodium nitrite solution. The enzyme activity was expressed as *µmol NO*^*−2*^ *gram fresh*.*weight*^*−1*^ *hour*^*−1*^.

### Statistical analysis

The data (Fresh and dry shoot/root weight, total protein content, and chlorophyll content, NR activity in leaf and root, Nitrite content and RWC) were analyzed by one-way analysis of variance (ANOVA) and the means were compared using the post hoc test of Tukey’s. All the data were computed using SPSS version 17. The means of three biological replicates were presented of all the traits.

### Protein Extraction

Total soluble proteins were extracted from three biological replicates of roots from control and stressed of each genotype separately. Approximately, 5 g of roots was ground with liquid nitrogen and homogenized in 10 mL of 0.5 M Tris-HCl, pH 7.5, lysis buffer containing 0.7 M sucrose, 50 mM EDTA, 0.1 M KCl, 10mM thiourea, 2mM PMSF and 2% v/v - mercaptoethanol. Then saturated phenols 10mL of Tris, pH 8 was added. After mixing for 30 min the phenolic phase was separated by centrifugation and rinsed with another 10 mL of lysis buffer. Protein was precipitated overnight at –20°C after adding 5 volumes of methanol containing 0.1 M ammonium acetate. The pellet recovered by centrifugation was rinsed with cold methanol and acetone, air-dried and re-suspended in 340 µl rehydration buffer. The protein content of samples (supernatants) were quantified by the method of Bradford **(Bradford 1976)** using bovine serum albumin (BSA) as a standard.

### 2DE-Gel Electrophoresis

Three biological and two technical replicates were performed form each genotype (CML49 and CML100) treatment and control plants. For first dimensional electrophoresis isoelectric focusing (IEF), was conducted by using 18cm strips (pH 3-10, linear gradient) in an *Ettan IPG phor IEF System* (*GE* Amersham) the total proteins supernatant from the roots tissue having 300 ug proteins in a buffer (9 M urea, 4% w/v CHAPS, 2% Triton X, 100 mM DTT and 2% v/v IPG buffer pH 3– 10 *(GE Healthcare)* was applied to the strips. The strips passively rehydrated for 2 hour and for next 10 hour active rehydration at 50V. The IEF was performed according to the following procedure: 250V-30min, 500V-30min, 10,000V gradient-3 hours, (until it reaches 70,000V hours) and finally 500V hold for 20min. After IEF, the strips were equilibrated for reduction and alkylation in equilibration buffer containing 6M urea, 1M Tris-HCl pH-8.8, 10% SDS, 30% Glycerol, 1% bromophenol blue. Initially the strips were treated with EB-1 (10mL EB+100mM Dithiothreitol) for 30 min, thereafter with EB-2 (EB+ 55mM Iodoacetamide) for 30 min. For second dimension the strips were loaded onto a 10% SDS-PAGE Gel *(Ettan Dalt Six Gel Electrophoresis Unit from GE)*, The molecular marker ladder of 250 KD-10 KD was applied on one side of the gels and strips were sealed with agros gel, after electrophoretic run at a currant of 10 mA per gel for 15 min, they adjusted to 30 mA till the dye reaches at the bottom of the plate. After 2DE, the gels were fixed in 10% methanol and 7% acetic acid for overnight and silver stained. The gels were scanned at 300 dpi to obtained digital images (TIFF files) using imagescanner *(Epson Expression 11000XL)* and spots detection was performed with Image Master 2D Platinum Software *(GE Healthcare)*. The spots were matched and detected by choosing a gel as the reference gel semi-automatically (manual correction and with default spot analysis setting in the software). Spots were normalized as the percentage of total spot volume that were present in all gels. The spot volume data was analyzed and Student t-test (p<0.05) was performed to verified statistically significant changes in spots. The molecular weight of the protein spots was calculated using the standard ladder applied to the gels, though pI of the protein spots determined with manufacturer’s instruction *(GE Healthcare)*. Spots were consider reproducible when they were also present in replicates gels. Differentially expressed protein spots with 1.5 or more fold variation in abundance were selected for mass spectrometry identification.

### Protein Digestion and Identification using MALDI-TOF/MS

Protein spots were excised manually from each set of gel, excised protein spots were de-stained in 400 μL of a 50% (v/v) acetonitrile, 25 mM NH4HCO3 solution for 30 min at room temperature. The procedure was repeated twice. The solution was discarded and 200 μL of pure acetonitrile was added for 5 min and dried under vacuum. The spots were reduced with 10mM DTT and alkylated with 50mM iodoacetamide (Bio-Rad). The proteins were then incubated for 30 min on ice in the presence of 10 μL of 20 ng/μL trypsin (Trypsin V5280, Promega), followed by 16 h at 37 °C. The peptides were extracted with 30 μL of a 5% trifluoroacetic acid solution. Then the extract was dried under vacuum and solubilized in 0.1% trifluoroacetic acid. The digested fragments were analyzed on MALDI-TOF analyzer (Autoflex II; Bruker Delatonics). Protein identification was performed using the Mascot software database (MSDB). The following parameters were used for database searches: taxonomy-Viridiplantae; enzyme-trypsin; fixed modifications-carbamidomethyl cysteine; variable modifications-oxidized methionine; peptide mass tolerance-0.25 Da; peptide charge-1 H+. Protein identifications were considered with a Mascot score >40 and sequence coverage of at least 25%.

## 3. Results

### Combined effects of drought, waterlogging stress and low-N stress on two maize genotypes

The various morpho-physiological parameters measured in two maize genotypes under combined abiotic stresses shows significant reduction in stressed plants compared to controls of both genotype, but the parameter decreases were more obvious in CML100 compared with CML49 **(Table 1)**. Similarly, drought stress significantly reduced leaf RWC greater in CML100 (54.33%) compared with CML49 (39.64%) respectively. The decrease in chlorophyll content was less obvious in CML100 (32.30%) and CML49 (34.25%) in drought stress. However, under waterlogging stress decreased in chlorophyll content in CML100 (44.56%) was much higher than CML49 (11.69%), thus under waterlogging stress CML49 shows steady chlorophyll content. Similarly, the difference in roots total protein content was less between CML49 (53.18%) and CML100 (52.04%). Nitrate Reductase, the main enzyme which plays a key role in nitrogen fixation responds to many environmental factors. Subsequently, low-N stress was applied to the plants (25% N of the recommended amount of urea), besides nutrients are less available under drought, or waterlogging conditions that may be low water availability, leaching or denitrification. Therefore, NR activity of roots was measured in both inbred plants after combined stresses and the results shows, Nitrate Reductase activity in CML49 was higher (2.019±0.236 *µmol NO*^*−2*^ *g fr*.*wt*^*−1*^ *hr*^*−1*^) relative to CML100 (0.673±0.180 *µmol NO*^*−2*^ *g fr*.*wt*^*−1*^ *hr*^*−1*^.), in CML49 the percent increase in NR activity was (>60%) high and significant than CML100 respectively. On the basis of physiological evaluation, we can say that CML49 shows more stable performance under combined stresses and could be categorized as ‘tolerant’. Whereas, CML100 was more stress sensitive and exhibited reduction in various morpho-physiological and biochemical parameters, hence consider as ‘sensitive’ inbred. Therefore, both genotypes have differential adaptability under combined stresses.

**Table 1.**
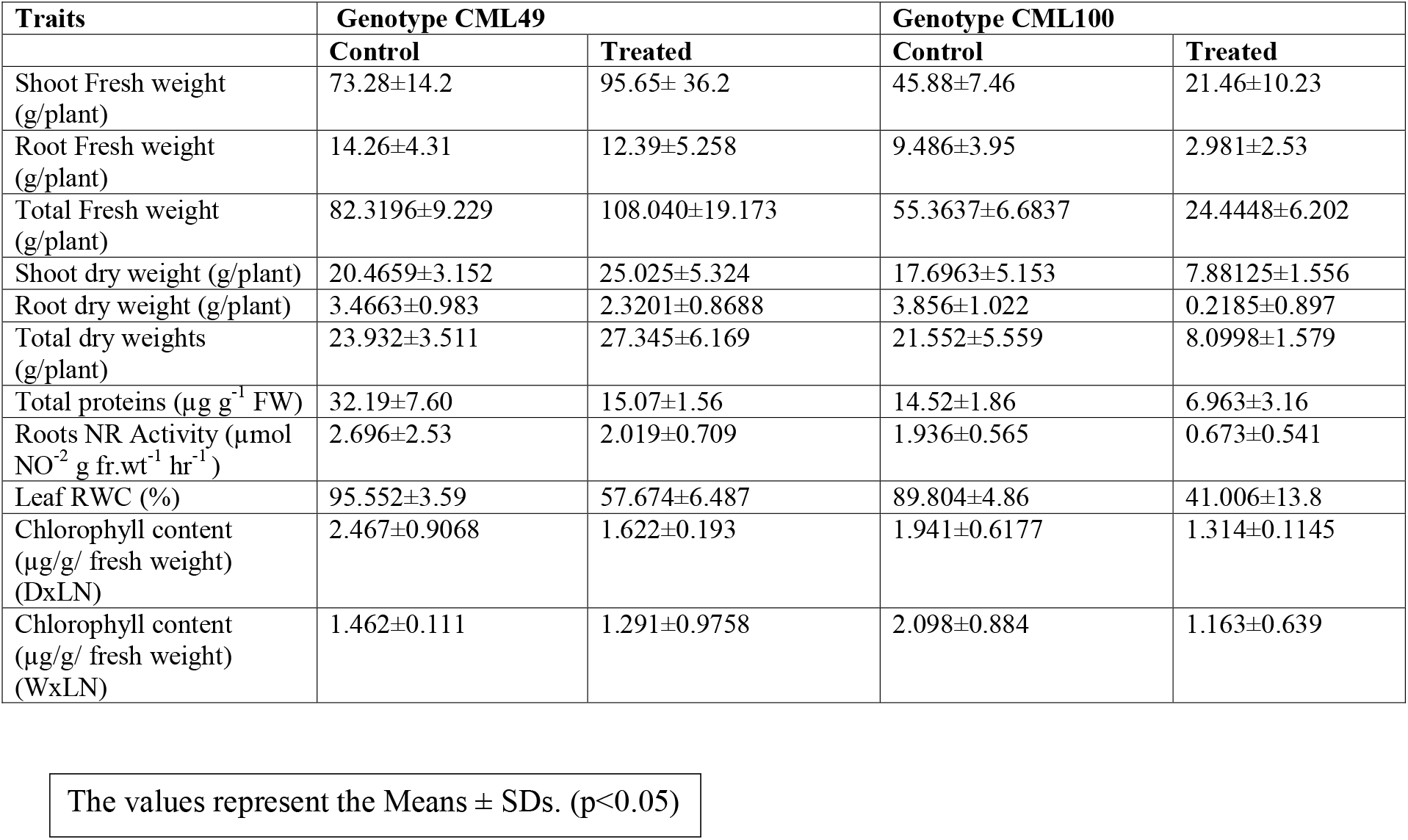
The various parameters measured in control and stressed plants of CML49 and CML100 inbred genotypes under combination of stresses.

### 2-DE gel analysis of roots proteome of two genotypes

The roots proteomic study was accomplish after exposing the two genotypes to combined stresses. The 2 DE gels of roots CML49 and CML100 were compared to their controls and differentially up-regulated protein spots were analyzed by MALDI-TOF by using Mascot database search for identification. Over 807 spots, were identified from each silver-stained gel of both genotype. The number of spots were higher in CML49 (418) compared to treated gels of CML100 (327). Among differentially up-regulated protein of spots of roots that exhibited quantitative changes greater than 1.5 fold variation in abundance, were selected for detailed analysis by MALDI-TOF. The total sixty three (63) spots up-regulated in CML49 **Fig 1**, whereas the up-regulated spots in CML100 was forty (40) **Fig 2**. The list of identified protein spots by Protein mass fingerprinting from both genotypes was shown **Table S1** (Supplementary) functional categorization of the proteins was done according to **M. Bevan et al (1998)**.

**Fig 1.**
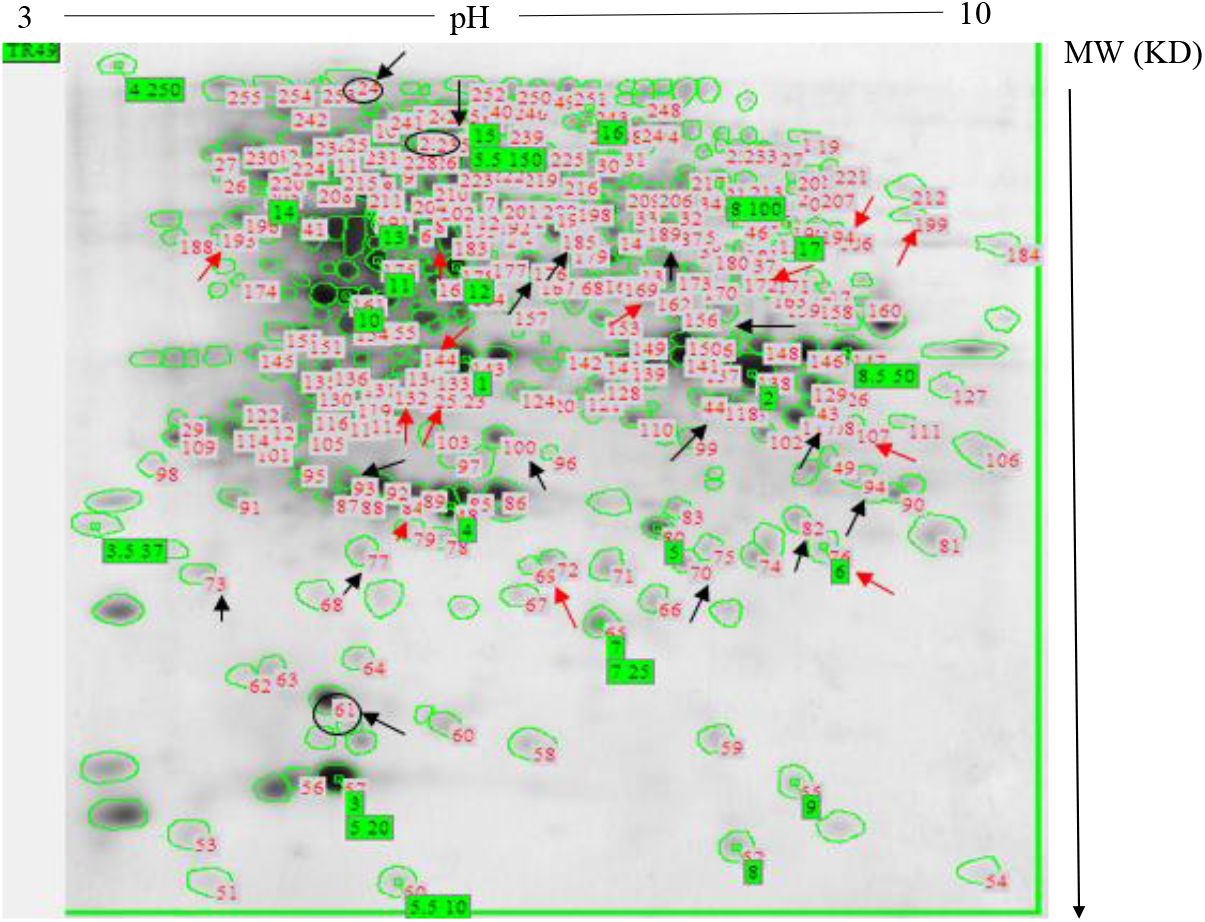
2-DE representative gel from CML49 (stressed roots) **2DE gel:** Fig 1 shows the representative gel of inbred CML49 roots. Black arrows-up regulated proteins spots. Red arrows-down regulated proteins spots. Spots with circle-represented the identified proteins spot by MALDI-TOF

**Fig 2.**
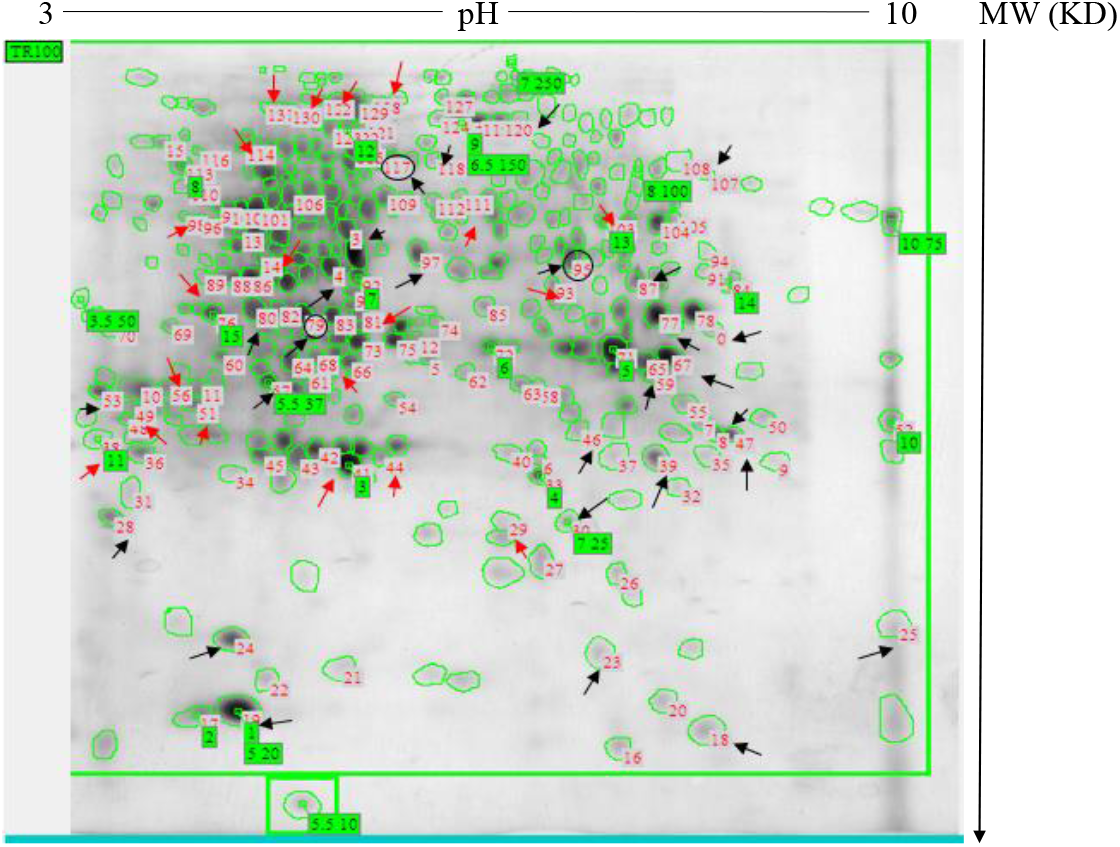
2-DE representative gel from CML100 (stressed roots) **2DE gel:** Fig 2 shows the representative gel of inbred CML100 roots. Black arrows-up regulated proteins spots. Red arrows-down regulated proteins spots. Spots with circle-represented the identified proteins spot by MALDI-TOF

## 4. Discussion

Plant growth and development depends on a vigorous root system, besides anchorage, roots plays key role in nutrient and water uptake. Our roots proteomics study shows the up-regulation of ‘Expansin protein’ in CML49 genotype. The activity of Expansin is associated with cell wall loosening in growing cells **(Lee et al. 2001)**, its localized expression in meristems and growth zones of roots, and also stem and leaf primordial **(Reinhardt et al. 1998)** was observed. Expansin, strongly regulated by water stress deficiency **(McQueen-Mason, 2004)**, similarly, in soybean, *GmEXPB2*, a gene that encodes a β-expansin (*EXPB*), also induced by mild water deficiency **(Guo et al 2011)** might involved in the response of the root system architecture. In maize roots, its activity increased by 4-fold (apical region) and by 2-fold (basal region) under low water potential, may be related to the higher expression of expansin encoding genes in the root apical zone, corresponding to maintaining elongation **(Wu et al. 1996; Wu et al. 2001)**. In contrast, other proteomic study showed in soybean seedlings expansin-like B1-like proteins were highly regulated in response to flooding stress and involved in cell wall metabolism **(Nakamura and Komatsu, 2013)**. Cyanoalanine hydratase (E.C. 4.2.1.65) is an enzyme involved in the cyanide detoxification process. One of the protein spot *Cynate hydrates (24)* up-regulated in the roots of CML49. Higher plants produced CN by six different mechanisms, including ethylene (ET) and camalexin biosynthesis, **(Yip and Yang 1988; Sanchez-Perez et al 2008)**. Higher plants maintaining cyanide homeostasis, through cyanide detoxification process that converts cyanide to β-cyanoalanine (β-Cas pathway), which is then converted to Asn, Asp, and ammonia by NIT4 class nitrilases, thus recycle of nitrogen within the plant (**Piotrowski 2008)**. Further, reports have shown that plants are capable of utilizing cyanide as a supplemental source of nitrogen (**Ebbs et al 2003; Siegien and Bogatek 2006)**. Higher NR activity of stressed roots and up-regulation of cynate hydrates proteins might attributed for tolerance to CML49 plants. The *tRNA (guanine-N (7) methyltransferase (223)* that catalyze the formation of N7-methylguanine at position 46 (m^7^G46) in tRNA. Also, several structurally different modified nucleosides are present in different tRNAs, at different position that have a common function to improved reading frame maintenance **(Urbonavieius et al 2001)**. Similarly, **Lang et al (2010)** associated it with drought tolerant as drought being a complex trait, many genes are involved it’s tolerance. Therefore, the up-regulated proteins played crucial role in tolerance mechanism of CML49 under various combined stress conditions. These results are well coherent with our leaf proteome study (Previous). On the contrary, CML100 shows higher activity of *Alcohol Dehydrogenase (95)* Alcohol dehydrogenase. (Fermentative enzyme that is highly conserved across species) several findings have confirmed that Alcohol dehydrogenase expression increased 6 h after flooding and decreased 24 h after draining water. Root apical meristem showed the strong induction of Adh2 expression in RNA and protein levels under flooding on the contrary, osmotic, cold, or drought stress has no expression of Adh2 genes **(Komatsu et al. 2011**). Further, at early vegetative stage, soya bean roots under waterlogging stress shows high up-regulation (9–16-fold) of alcohol dehydrogenase (Adh) protein and may involve in continuing glycolysis **(Iftekhar Alam et al. 2010)**. Later, **Ismond et al (2013)** showed the role of ADH in recycling of NADH to NAD+, for the continuation of glycolysis pathway and is the only energy source under oxygen deprivation condition. However, it had no effect on flooding survival **(Ismond et al 2003)**. The induction of *Caffeic acid 3-O-methyltranferase (79)*, protein in CML100 roots and similar induction (CCoAOMT enzyme) was observed under salinity stress in the roots of Arabidopsis and rice **(Salekdeh et al 2002; Lee et al 2008)**. The up regulation of *Sucrose Phosphate Synthase (117)*, shows that in photosynthetic and non-photosynthetic tissues SPS is regulated through metabolites and protein phosphorylation **(Reimholz et al 1994)**. Likewise in wheat, SPP gene family constitutively expressed in source (leaves and germinating seeds) and sink tissues (developing seeds and roots) **(Lunn 2003)**. Recently, **Gloria et al (2017)** observed the expression of sps gene; AtSPS1F, AtSPS2F and AtSPS3F in the columella roots of A thaliana plants hence, the findings support that sucrose synthesis can occurs in the columella cells, and transcriptional analysis shows AtSPS2F and AtSPS4F activated in response to osmotic stress. Similar studies by **Kaur et al (2007)** under water deficit condition in tolerant cultivar and sensitive cultivar of wheat seedlings exhibited the Sucrose synthase activity was lower in the shoots and roots of stressed seedlings of tolerant cultivar. These findings in agreement of our results and suggest the possible role of SPS protein of CML100 plant.

## 5. Conclusion

The Physio-biochemical studies shows the response two genotypes differed in various combined stresses. The CML49 was tolerant and acclimatized to the various abiotic stresses, whereas, CML100 shows sensitive behavior towards multiple stresses. The present study shows up-regulated proteins of roots of CML49 plants under multiple stress conditions are involved in cell wall remodeling, cyanide detoxification, and drought tolerant mechanism. Thus, roots proteins were in alliance with leaf proteome of tolerant inbred. In same way, up-regulated proteins of CML100 plants involved in root primary and secondary metabolism to protect against the stresses and biomarker of anoxia stress, the result was coherent with leaf proteome. In our previous work we have shown that **(Suphia R, 2019)**, up-regulated leaves proteins of tolerant and susceptible maize plants in response to multiple abiotic stresses were involved in enhancing tolerance mechanism of tolerant inbred and up-regulated proteins of susceptible inbred help to survive the stressful conditions. Therefore, the study provides the comprehensive analysis of tolerance mechanism that can be further utilized to develop the climate resilient crop varieties.

## Supporting information

Identified Spots of tolerant and sensitive genotypes by MALDI-TOF

## DECLARATIONS

## Acknowledgments/ Grant received

I acknowledge the grant received from Department of Science and Technology, Ministry of Science, Government of India, under the WOS-A Scheme.

***Reference No: SR/WOS-A/LS-284/2013***

## Availability of data and material

***Table S1:*** The following files below contains the detailed roots Proteomics data (Supplementary data files)

## Authors Contributions

Suphia Rafique has contributed to the study conception and design. Material preparation, data collection and analysis and interpretation, writing the manuscript. The Mentor of the Project is [Professor M Z Abdin]. The first draft of the manuscript was written by [Suphia Rafique] and all authors read and approved the final manuscript

## Abstract

### Purpose

Abiotic stresses are the major threat to crops regardless of their nature, duration, and frequency, their occurrence either singly, and or combined. Maize is the most important crop largely grown in the tropical region in the summer rainy season, often faces a stress combination of drought and waterlogging. We previously showed under multiple stresses up-regulated leaf proteins of maize plants are involved to enhance the tolerance mechanism of tolerant genotype and up-regulated proteins in susceptible genotypes help to survive the stressful condition.

### Method

To understand the response of two maize genotypes to various combined stresses, physio-biochemical analysis was done. Further, to understand the molecular aspects roots proteomics study was achieved using the 2DE technique.

### Results

The physio-biochemical analysis of the two inbred shows a difference in adaptability to the various combined stresses, CML49 shows tolerance and CML100 was sensitive towards the various stresses. The root proteome shows the up-regulated proteins of CML49 genotype are involved in enhancing the N content, cell wall remodeling, and acclimatization during the stresses. Up-regulated proteins of CML100 genotype were stressed marker of roots primary and secondary metabolism.

### Conclusion

However, the root proteome of both genotypes correlates with the leaf proteome (previous). Therefore, the present study and our previous results provide comprehensive insight into the molecular mechanisms of tolerance in combined abiotic stresses of maize plants.

## REFERNCES

Atkinson N J, Lilley C J, and Urwin P E (2013) Identification of genes involved in the response to simultaneouss biotic and abiotic stress Plant Physiol 162: 2028–2041

Bradford M M, (1976) A rapid and sensitive method for the quantitation of microgram quantities of protein utilizing the principle of protein-dye binding. Anal Biochemistry 721 (2): 248–254

Bevan M, et al (1998) Analysis of 19 Mb of contiguous sequence from chromosome 4 of Arabidopsis thaliana Nature 391: 485–488

Choudhary A Pandey P and Senthil-Kumar M (2016) “Tailored responses to simultaneous drought stress and pathogen infection in plants” in Drought Stress Tolerance in Plants Vol 1 eds M A Hossain S H Wani S Bhattacharjee D J Burritt and L- SP Tran (Cham: Springer International Publishing) 427–438

Ebbs S J Bushey S Poston D Kosma M Samiotakis D Dzombak (2003) Transport and metabolism of free cyanide and iron cyanide complexes by willow. Plant Cell Environ, 26: 1467–1478

Guo W, Zhao J Li, X Qin L, Yan X, Liao H (2011) A soybean b expansin gene GmEXPB2 intrinsically involved in root system architecture responses to abiotic stresses. Plant J 66(3):541–552

Solís-Guzmán Marí Gloria, Argüello-Astorga G, López-Bucio José Ruiz-Herrera. LeóFrancisco López-Meza JE, Sánchez-Calderón L, Carreón-Abud Yazmí, Martínez-Trujillo M Arabidopsis thaliana sucrose phosphate synthase (sps) genes are expressed differentially in organs and tissues and their transcription is regulated by osmotic stress Gene Expression Patterns 2017

Hageman R H, and Hucklesby D P (1971) Nitrate reductase from higher plants Meth Enzymol 23: 491–503

Iftekhar Alam, Dong-Gi Lee, Kyung-Hee Kim, Choong-Hoon park, Shamima Akhtar, Sharmin Hyoshin, Lee Ki-Won O H, Byung-Wook Yun and Byung-Hyun (2010) Proteome analysis of soybean roots under waterlogging stress at an early vegetative stage Journal of Bioscience 35: 1 49–62

Ismond K P, Dolferus R dePauw M, Dennis E S, and Good A G (2013) Enhanced low oxygen survival in Arabidopsis through increased metabolic flux in the fermentative pathway. Plant Physiol 132: 1292–1302

Ismond K P, Dolferus R de Pauw M, Dennis ES, Good AG (2003) Enhanced low oxygen survival in Arabidopsis through increased metabolic flux in the fermentative pathway. Plant Physiol 132 (3): 1292–302

Kaur K, Gupta A K, Kaur N (2007) Effect of water deficit on carbohydrate status and enzymes of carbohydrate metabolism in seedlings of wheat cultivars. Indian Journal of Biochemistry and Biophysics 44: 223–230

Klára Kosová, Pavel Vítámvás, Milan O Urban, Ilja T Prášil, and Jenny Renaut (2018) Plant Abiotic Stress Proteomics The Major Factors Determining Alterations in Cellular Proteome. Frontiers in Plant Science 122: 9 1-22–322

Komatsu S, Deschamps Thibaut, Susumu Hiraga, Mikio Kato Mitsuru, Chiba Akiko Hashiguchi Makoto Tougou, Satoshi Shimamura, Hiroshi Yasue (2011) Characterization of a novel flooding stress-responsive alcohol dehydrogenase expressed in soybean roots. Plant Molecular Biology 77: 309

Lang Nguyen, Thi Nguyen, Quang Cao, Binh Chau Thanh, Nha Bui, Chi Buu (2010) A candidate gene response to drought stress condition in rice (Oryza sativa L). Omonrice 17: 105–113

Lee Y Choi D Kende H: Expansins: ever-expanding numbers and functions Curr Opin Plant Biol 2001 4:527–532

Lee YJ, Kim BG, Chong Y et al (2008) Cation dependent O-methyltransferases from rice. Planta 227: 641–647

Lunn J E (2003) Sucrose-phosphatase gene families in plants. Gene 303: 187–196

McQueen-Mason S Jones L (2004) A role for expansins in dehydration and rehydration of the resurrection plant Craterostigma plantagineum. FEBS Lett 559: 61–65

Nanjo Y Nakamura, T Komatsu S (2013) Identification of indicator proteins associated with flooding injury in soybean seedlings using label-free quantitative proteomics. J Proteome Res 12: 4758–4798

Oh M and Komatsu S (2015) Characterization of proteins in soybean roots under flooding and drought stresses. J Proteomics 114: 161–181

Pandey P Irulappan, V Bagavathiannan MV, and Senthil-Kumar M (2017) Impact of Combined Abiotic and Biotic Stresses on Plant Growth and Avenues for Crop Improvement by Exploiting Physio-morphological Traits. Front Plant Sci 8:537

Peng Z, Wang M, Li F L H Li C, and Xia G (2009) A proteomic study of the response to salinity and drought stress in an introgression strain of bread wheat. Mol Cell Proteomics 8: 2676–2686

Piotrowski M (2008) Primary or secondary? Versatile nitrilases in plant metabolism. Phytochemistry 69: 2655–2667

Prasch C M, and Sonnewald U (2013) Simultaneous application of heat drought and virus to Arabidopsis plants reveals significant shifts in ignaling networks. Plant Physiol 162: 1849–1866

Ramu V S, Paramanantham A, Ramegowda V, Mohan-Raju B, Udayakumar M, and Senthil-Kumar M (2016) Transcriptome analysis of sunflower genotypes with contrasting oxidative stress tolerance reveals individual- and combined-biotic and abiotic stress tolerance mechanisms. PLoS ONE 11:e0157522

Rathore T R, MZK Warsi, JE Lothrop, NN Singh (1996) Production of maize under excess soil moisture waterlogging conditions pp 56-63 In: 6th Asian Regional Maize Workshop 10-12 Feb 1996 PAU Ludhiana

Ralph Reimholz, Peter Geigenberger, Mark Stitt (1994) Sucrose-phosphate synthase is regulated via metabolites phosphorylation in potato tubers in a manner analogous to the enzyme in leaves. Planta 192: 480–488

Reinhardt D, Wittwer F, Mandel T, Kuhlemeier C (198) Localized upregulation of a new expansin gene predicts the site of leaf formation in the tomato meristem. Plant Cell 10:1427–1437

Rizhsky L, Liang H, Shuman J, Shulaev V, Davletova S, Mittler R (2004) When defense pathways collide The response of Arabidopsis to a combination of drought and heat stress. Plant Physiology. 134: 1683–1696.

Rollins J A, Habte E, Templer S E, Colby T, Schmidt J and von Korff M (2013) Leaf proteome alterations in the context of physiological and morphological responses to drought and heat stress in barley (Hordeumvulgare L). J Exp Bot 64: 3201–3212

Salekdeh GH, Siopongco J, Wade LJ, Ghareyazie B, Bennett J (2002) Proteomic analysis of rice leaves during drought stress and recovery. Proteomics 2: 1131–1145

Sanchez-Perez R, Jorgensen K, Olsen C E, Dicenta F, Moller B L (2008) Bitterness in almonds Plant Physiol 146(3):1040–1052

Suphia Rafique (2019) Differential expression of leaf proteomes of tolerant and susceptible maize (Zea mays L) genotypes in response to multiple stresses. Biochem Cell Biol 97: 581–588 dxdoiorg/101139/bcb-2018-0338

Suzuki N, Rivero R M, Shulaev V, Blumwald E and Mittler R (2014) Abiotic and biotic stress combinations. New Phytol 203: 32–43

Siegien I. and R Bogatek (2006) Cyanide action in plants: from toxic to regulatory. Acta Physiol Plant 28: 483–497

Urbonavieius J, Qian Q, Durand J M B, Hagervall T G, and Bjork G R (2001) Improvement of reading frame maintenance is a common function for several tRNA modification. EMBO J 20: 4863–4873

Vile D, Pervent M, Belluau M, Vasseur F Bresson, J Muller B, et al (2012) Arabidopsis growth under prolonged high temperature and water deficit: independent or interactive effects? Plant Cell Environ 35: 702–718

Wu Y, Sharp RE, Durachko DM, Cosgrove DJ (1996) Growth maintenance of the maize primary root at low water potentials involves increases in cell-wall extension properties expansin activity and wall susceptibility to expansins. Plant Physiol 111: 765–772

Wu Y, Thorne ET, Sharp RE, Cosgrove DJ (2001) Modification of expansin transcript levels in the maize primary root at low water potentials. Plant Physiol 126: 1471–1479

Yip W K, Yang S F 91998) Cyanide metabolism in relation to ethylene production in plant tissues. Plant Physiol 88(2):473–476

Zaidi PH, Mamata Yadav, DK Singh, and RP Singh (2008) The relationship between drought and excess moisture tolerance in tropical maize Zea mays L. Aust J Crop Sci 13: 78–96

Wen W Li K Alseekh S Omranian N Zhao L Zhou Y et al Genetic determinants of the network of primary metabolism and their relationships to plant performance in a maize recombinant inbred line population Plant Cell 2015 27: 1839–1856

